# Structural Characterization of L-Galactose Dehydrogenase: A Key Enzyme for Vitamin C Biosynthesis

**DOI:** 10.1101/2022.04.15.488469

**Authors:** Jhon A. Vargas, Diego A. Leonardo, Humberto D’Muniz Pereira, Adriana R. Lopes, Hicler N. Rodriguez, Marianela Cobos, Jorge L. Marapara, Juan C. Castro, Richard C. Garratt

**Author notes:** Corresponding author: R. C. Garratt, São Carlos Institute of Physics, University of São Paulo, Avenida João Dagnone 1100, São Carlos, SP 13563-120, Brazil. These authors contributed equally (Jhon A. Vargas, Diego A. Leonardo).

## Abstract

In plants, it is well-known that ascorbic acid (vitamin C) can be synthesized via multiple metabolic pathways but there is still much to be learnt concerning their integration and control mechanisms. Furthermore, the structural biology of the component enzymes has been poorly exploited. Here we describe the first crystal structure for an L-galactose dehydrogenase (SoGDH from spinach), from the D-mannose/L-galactose (Smirnoff Wheeler) pathway which converts L-galactose into L-galactono-1,4-lactone. The kinetic parameters for the enzyme are similar to those from its homologue from camu-camu, a super-accumulator of vitamin C found in the Peruvian amazon. Both enzymes are monomers in solution, have a pH optimum of 7 and their activity is largely unaffected by high concentrations of ascorbic acid, suggesting the absence of a feedback mechanism acting via GDH. Previous reports may have been influenced by changes of the pH of the reaction medium as a function of ascorbic acid concentration. The structure of SoGDH is dominated by a (β/α)_8_ barrel closely related to aldehyde-keto reductases (AKRs). The structure bound to NAD^+^ shows that the lack of Arg279 justifies its preference for NAD^+^ over NADP^+^, as employed by many AKRs. This favours the oxidation reaction which ultimately leads to ascorbic acid accumulation. When compared with other AKRs, residue substitutions at the C-terminal end of the barrel (Tyr185, Tyr61, Ser59 and Asp128) can be identified to be likely determinants of substrate specificity. The present work contributes towards a more comprehensive understanding of structure-function relationships in the enzymes involved in vitamin C synthesis.

## Introduction

L-Ascorbic Acid (ascorbate or AsA), commonly called vitamin C, is an essential biomolecule with key pleiotropic functions in both animals and plants. In animals, in addition to its free radical-scavenging activity, AsA functions as a cofactor for hydroxylases, mono-oxygenases and dioxygenases (Englard and Seifter, 1986; Fenech et al., 2019). Furthermore, emerging evidence suggests that AsA participates in the regulation of relevant epigenomic processes such as the modulation of both DNA methylation and histone chemical modification (Nur et al., 2021; Young et al., 2015). Similarly, in plants AsA has a strong antioxidant activity, is a cofactor for mono- and dioxygenases (including enzymes for the biosynthesis of phytohormones and anthocyanins), acts as a photoprotectant and controls plant growth through cell division and expansion (Smirnoff, 2000; Smirnoff and Wheeler, 2000).

In plants, there are multiple biosynthetic routes for the production of AsA (Caruso et al., 2021; Castro et al., 2015; Liao et al., 2021; Valpuesta and Botella, 2004). Of these, the best characterized is the D-mannose/L-galactose (or Smirnoff-Wheeler) pathway (Wheeler et al., 1998, Figure S1). Despite this considerable effort, structural information concerning its component enzymes is still very limited. One of these is L-galactose dehydrogenase (L-GDH, EC 1.1.1.316), which catalyzes the oxidation of L-galactose (L-gal) at position C1 to produce L-galactono-1,4-lactone in the penultimate step of the synthesis of AsA. L-GDH was one of the first enzymes of the pathway to be properly studied, being recognized as one of the most important in the regulation of AsA biosynthesis (Gatzek et al., 2002; Laing et al., 2004; Mieda et al., 2004; Wheeler et al., 1998).

Biochemical studies of L-GDH, either purified from leaves or via heterologous expression, have been carried out on different species including *Pisum sativum* (*Ps*GDH), *Arabidopsis thaliana* (*At*GDH), *Spinacia oleracea* (*So*GDH) and *Actidinia deliciosa* (*Ad*GDH) (Gatzek et al., 2002; Laing et al., 2004; Mieda et al., 2004). L-GDH shows a high affinity and selectivity for L-gal with *Km* ranging from 0.08-0.43 mM, as well as a preference for NAD^+^ rather than NADP^+^ as its cofactor (Gatzek et al., 2002; Laing et al., 2004; Mieda et al., 2004). Furthermore, AsA has been shown to slowly and irreversibly inactivate the enzyme or act as a feedback inhibitor through competitive inhibition with the substrate. This suggests that L-GDH may play a significant role in the regulation of AsA biosynthesis in plants (Laing et al., 2004; Mieda et al., 2004). However, this potential regulatory mechanism for the biosynthesis of AsA needs to be further clarified at the molecular level. Despite the extensive knowledge about the biochemical characteristics of L-GDH, together with two entries for the enzyme from rice (*Oryza sativa, Os*GDH) in the PDB (7EZL and 7EZI), its structural characteristics have yet to be described in the literature.

*Myrciaria dubia* (Kunth) McVaugh, also known as camu-camu, is a fruit, native to the Amazon rain forest. Camu-camu is characterized by its high concentration of vitamin C in fruits, which varies between 0.96 – 2.99 g/100 g of pulp (Castro et al., 2018). By comparison, species which are considered to be references for vitamin C content, such as oranges or kiwis, have concentrations of only 0.05 g and 0.085 g/100 g of pulp, respectively (Locato et al., 2013). This characteristic allows us to speculate that the enzymes from camu-camu may have potential biotechnological applications in the artificial production of vitamin C.

Camu-camu presents five metabolic pathways for vitamin C biosynthesis, including the D-mannose/L-galactose pathway (Castro et al., 2015). Here, we purify the recombinant L-galactose dehydrogenase from *Myrciaria dubia* (*Md*GDH) and compare its biochemical properties with that from spinach (*So*GDH). In so doing we shed new light on the reported role of AsA in feedback inhibition. Finally, for the first time, we describe the crystal structure of the spinach enzyme in both its apo and NAD^+^ bound forms (*So*GDH), allowing for the elucidation of the cofactor-induced conformational change and the identification of residues potentially important for substrate specificity.

## Results and Discussion

### The oligomeric state of L-galactose dehydrogenase

Heterologously expressed camu-camu *Md*GDH and spinach *So*GDH showed a single peak on size exclusion chromatography (SEC) and a predominant band on SDS-PAGE (figure S2A). Consistent with this result, the size exclusion chromatography coupled with multi-angle light scattering (SEC-MALS) profiles showed a predominant monodisperse peak, corresponding to the expected molecular mass of the monomeric state for both recombinant enzymes (figure S2B). In contrast, in some previous investigations, purified L-GDH exhibited variation in the oligomeric state, depending on the plant species and source employed. For example, *Ps*GDH purified from *Pisum sativum* (pea) embryonic axis was reported to be a tetrameric protein (Gatzek et al., 2002). On the other hand, *So*GDH and *Ad*GDH, purified from the leaves of *Spinacia oleracea* (spinach) and *Actinidia deliciosa* (kiwifruit), were described as being homodimers and monomers, respectively (Laing et al., 2004; Mieda et al., 2004). Similarly, the recombinant enzymes from *Arabidosis thaliana* (*At*GDH) and *Oryza sativa* (*Os*GDH) were reported to be homodimeric and monomeric, respectively (Gatzek et al., 2002; Momma and Fujimoto, 2013). Although there are differences in the techniques used for protein purification from natural or heterologous sources, this would not normally be expected to affect the oligomeric state of the enzyme. Moreover, the investigations that reported the enzyme to form dimers or tetramers provided relatively limited data to substantiate these claims (Gatzek et al., 2002; Mieda et al., 2004). By contrast, the studies that reported the monomeric state for this enzyme used size exclusion chromatographic analysis (Laing et al., 2004; Momma and Fujimoto, 2013).

The monomeric state of both L-GDHs studied here, which appears to be unambiguous from our SEC-MALS results, is also consistent with the inclusion of L-GDH as part of the aldo-keto reductases (AKR) protein superfamily (Gatzek et al., 2002). This is comprised of 16 different families that are distributed across all phyla. These enzymes catalyze oxidation-reduction reactions on carbonyl substrates (eg., sugar aldehydes, keto-steroids, quinones, etc.) and usually are monomers whose fold is dominated by a TIM-barrel motif (β/α)_8_. They are further characterized by a conserved cofactor binding region for NAD(P)H, variable loop structures that define substrate specificity, and a catalytic tetrad (Hyndman et al., 2003; Jez et al., 1997; Penning, 2015). Of the 16 families of AKRs (Penning, 2015), only three dimeric AKRs have been reported: a rat liver aflatoxin dialdehyde reductase (AKR7A1), the *Candida tenuis* NAD(P)H dependent xylose reductase (AKR2B5), and the *Escherichia coli* tyrosine auxotrophy suppressor protein (Kavanagh et al., 2003; Kozma et al., 2002; Obmolova et al., 2003). The crystal structures of these proteins shows a conserved contact interface for dimer formation, which is absent from the enzymes described here. This will be discussed later when describing the crystal structure of *So*GDH and adds weight to the expectation that L-GDH is, in general, a monomeric enzyme.

### High concentrations of AsA induce inhibition of L-GDH by pH changes

Camu-camu *Md*GDH shows Michaelis-Menten-like kinetics using L-gal as substrate, with an optimal pH value of 7.0 and *Km* of 0.21 mM (figure 1A). According to recent reports (Gatzek et al., 2002; Laing et al., 2004; Mieda et al., 2004), the *Km* value for L-GDH from different plant species varies from 0.08 to 0.43 mM, and consequently, *Md*GDH can be considered to be an enzyme with a medium affinity for the L-gal substrate (table 1). Furthermore, *Md*GDH has a turnover number (*kcat*) of 4.26 s^-1^ and a catalytic efficiency (*kcat*/*Km*) of 20.67 mM^-1^/s^-1^. These kinetic parameters are being reported here for the first time for any species so far studied. The spinach recombinant enzyme (*So*GDH) presented a *Km* of 0.13 mM (figure 1D), close to that reported for *So*GDH purified from spinach leaves (0.12 mM, Mieda et al., 2004). By contrast, however, we observed an optimal pH of 7.0 (similar to *Md*GDH) while Mieda et al. reported a value of 9.25. For *So*GDH we observed a *kcat* of 1.2 s^-1^ and a catalytic efficiency of 9.1 mM^-1^/s^-1^ (table 1). Overall, the two enzymes show kinetic parameters which are broadly similar and well within an order of magnitude of one another for all values. The discrepancy in the reported values for the optimal pH of *So*GDH is difficult to explain at this moment but it is of note that the report by Mieda et al (2004) provides no experimental detail on how the value of 9.25 was determined. Furthermore, the authors only cite the use of Tris buffer which would be inappropriate for investigating a wide pH range. Finally, despite quoting pH 9.25 as the optimal pH for spinach GDH, all of their kinetic studies were performed in 100 mM Tris buffer at pH 7.5.

**Table 1.** Kinetics parameters for plant L-GDHs.

| Enzymes | $K_m$<br>(mM) | $K_{cat}$<br>($S^{-1}$) | $K_{cat}/K_m$<br>($mM^{-1}.S^{-1}$) | $K_i$<br>(mM) | optimal<br>pH |
| --- | --- | --- | --- | --- | --- |
| **Camu-Camu *MdGDH*<sup>€</sup>** | 0,206 | 4.261 | 20,67 | 1,2 | 7,0 |
| **Spinach *SoGDH*<sup>€</sup>** | 0.128 | 1,158 | 9,06 | 0,5 | 7,0 |
| Arabidopsis *AtGDH*<sup>£</sup> | 0.085 | N/D | N/D | N/D | 8,5-9 |
| Pea *PsGDH*<sup>£</sup> | 0.430 | N/D | N/D | N/D | 8,5-9 |
| Kiwi *AdGDH*<sup>¤</sup> | 0.300 | N/D | N/D | 0,5 | 9,0 |
| Spinach *SoGDH*<sup>*</sup> | 0.116 | N/D | N/D | 0,1 | 9,25 |

<sup>€</sup>L-GDH parameters according to this study.
<sup>£</sup>L-GDH parameters according to Gatzek et al, 2002
<sup>¤</sup>L-GDH parameters according to Laing et al, 2004
<sup>\*</sup>L-GDH parameters according to Mieda et al, 2004.

**Figure 1.**
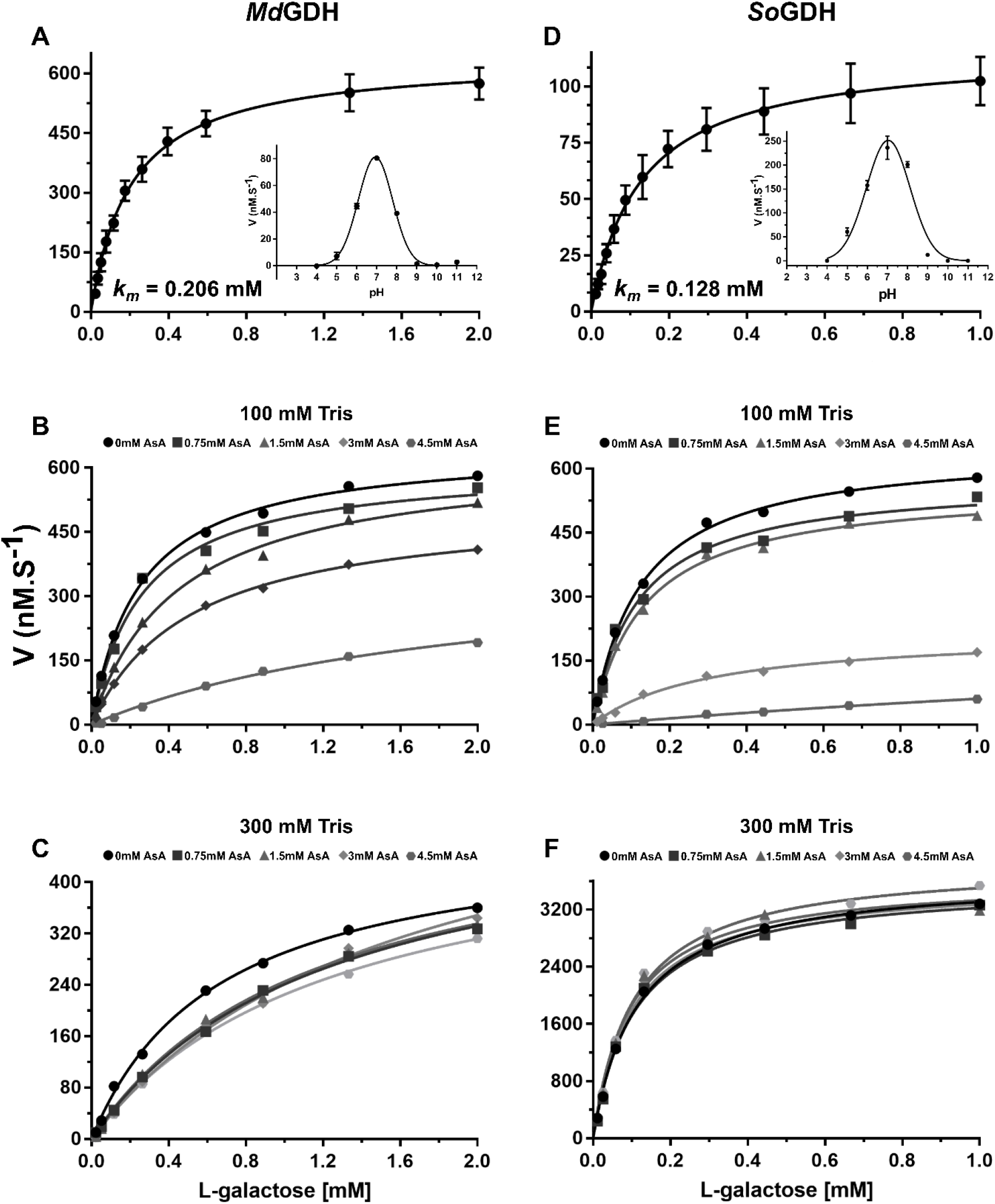
Kinetic properties of camu-camu *Md*GDH and spinach *So*GDH in the presence of excess NAD^+^ (200 μM). Michaelis-Menten-like type kinetics were observed for *Md*GDH (A) and *So*GDH (D) with *Km* values of 0.206 and 0.128. mM, respectively. Optimum pH curves are shown as insets. The inhibition of *Md*GDH and *So*GDH by AsA in either 100 mM Tris-HCl buffer (B and E) or 300 mM Tris-HCl (C and F) are shown. The inhibitory effect of AsA drops off drastically in the higher concentration buffer.

The study of Mieda *et al*. (2004) also suggested feedback regulation of L-GDH by AsA, the end-product of the D-mannose/L-galactose biosynthetic pathway, by competitive inhibition. According to this report, AsA at 1 mM inhibits up to 41% of *So*GDH activity with a *Ki* of 0.13 mM and L-GDH purified from *Arabidopsis thaliana* was similarly inhibited by AsA (Mieda et al., 2004). However, a more detailed analysis of the inhibitory effect of AsA on the enzyme from kiwi fruit (*Ad*GDH) showed that the inactivation is essentially slow and irreversible, and can be attributed to oxidative damage of a key amino acid residue in the active site (Laing et al., 2004).

Camu-camu is a plant that stores high concentrations of AsA (2g/100g pulp) in its fruits (Castro et al., 2018) and consequently *Md*GDH could potentially be refractory to feedback regulation by AsA. In order to test this hypothesis, we conducted inhibition assays of *Md*GDH and *So*GDH with AsA concentrations up to 4.5 mM (figure 1B). AsA at 1.5 mM, 3 mM, and 4.5 mM inhibited the activity of *Md*GDH by 53%, 71% and 79%, respectively (*Ki* = 1.2 mM). At the same AsA concentrations the activity of *So*GDH was inhibited by 82%, 89% and 92%, respectively (*K*i= 0.5 mM) (figure 1E), indicating that camu-camu *Md*GDH has a slightly higher tolerance to inhibition by AsA than does spinach *So*GDH and kiwi *Ad*GDH.

Although AsA is only a weak acid, the concentrations used in the inhibition assays could lead to significant alterations to the pH of the reaction buffer (100 mM Tris-HCl, pH 7.0). Considering this effect and knowing that changes in pH alter the activity of L-GDH (figure 1), we performed pH measurements of the reaction buffer with each of the AsA concentrations used in the assay. We observed that on increasing the AsA concentration, the pH of the reaction solution decreased. For example, using a reaction solution with Tris-HCl pH 7.0 at 100 mM, the pH droped to 5.9 in the presence of 4.5 mM AsA (table S1). The classical bell-shaped pH dependence curves shown in the inset to figures 1A and 1D indicate that this pH drop of > 1 pH unit would have a significant impact on enzyme activity, of the order we observed experimentally. This suggests that the inhibition observed in both camu-camu and spinach L-GDH could be the effect of pH and not a genuine feedback inhibition by AsA, the pathwayś final product. In order to investigate this further, inhibition assays were performed in the presence of a stronger buffer (300 mM Tris-HCl) where the pH of the reaction buffer is not drastically affected by high concentrations of AsA (Table S1). Under these conditions, AsA showed no inhibitory effect on the activity of *So*GDH (figure 1F). However, in the case of *Md*GDH mild inhibition persisted with an estimated *Ki* of 2.1 mM. This effect may be related to the reaction buffer itself, since when comparing the *Km* values of the enzyme in the absence of AsA in the 100- and 300-mM buffers, it varies from 0.206 mM to 0.650 mM, respectively. Nevertheless, the effect produced by AsA on the enzyme activity appears to be largely due to a drop in pH in the case of both enzymes. It is worth noting that Mieda *et al*. (2004) also performed their study using Tris buffer at only 100mM concentration. Our data call into question the conclusions of the study of Mieda et al. (2004) who suggest negative feedback inhibition by the final product of the pathway (AsA) as a potential regulatory mechanism of physiological relevance to plants. We find no evidence for this in the present study.

### Overall description of the structure of L-GDH

To date, no crystal structures for L-GDHs have been described in the literature. In order to fill this gap in our current knowledge we embarked upon a crystallization campaign for both of the enzymes of interest to the current study. From crystallization assays performed on camu-camu *Md*GDH and spinach *So*GDH, X-ray diffraction quality crystals were obtained for *So*GDH in both its apo and holo (NAD^+^ bound) forms. Diffraction data for *So*GDH and *So*GDH-NAD^+^ were collected, and the structures refined to 1.40 and 1.75 Å resolution, respectively. The good quality of the models is reflected in the values of Rwork and Rfree, as well as in their stereochemistry. *So*GDH has a final Rwork = 19.42% and Rfree = 20.95%, with 97.76% of amino acids in the most favoured regions of Ramachandran space. In the case of *So*GDH-NAD^+^ the equivalent values are =18.37%, 21.48%, and 98.08% respectively (table S2 summarizes the full data collection and refinement statistics).

A preliminary X-ray analysis of *Os*GDH from rice (*Oryza sativa*) was reported in 2013 but the structure remained unsolved due to difficulties with molecular replacement (Momma and Fujimoto, 2013). Nevertheless, in 2021 the same authors deposited two structures for the enzyme, one in its apo form at 1.2 Å (*Os*GDH [PDB accession: 7EZI]) and one in the form of a complex with the cofactor NAD^+^ at 1.8 Å (*Os*GDH-NAD^+^ [PDB accession: 7EZL]). However, thus far, no publication describing the details of these first structures has appeared in the literature. At the time when the structures reported here were solved, the homologous structures from rice were not yet available and initially we had similar difficulties in encountering a correct molecular replacement solution. However, the structure of the apo enzyme (*So*GDH) was finally solved using MoRDa, a pipeline for automated molecular replacement based on a protein domain database derived from the PDB (Vagin and Lebedev, 2015). The structure of the cofactor-bound complex (*So*GDH-NAD^+^) was solved by molecular replacement using the apo structure as the search model with the program *Phaser* in the *Phenix* package (Adams et al., 2010).

The crystal structure of *So*GDH has one molecule in the asymmetric unit with a (β/α)_8_-barrel fold, as observed in the AKR superfamily (figure 2A) (Jez et al., 1997). It has 8 parallel β-strands which are alternated with 8 α-helices which run antiparallel relative to the strands, forming the classical barrel-type fold. At the N-terminus, is an additional β-hairpin (β1 and β2) that forms the bottom of the barrel. The (β/α)_8_-barrel is conserved in the AKR superfamily, however the loops connecting the strands and helices, as well as the presence of external helices to the barrel, can vary widely and are important for the classification of AKRs into the 16 different families (Penning, 2015). The Loops A (β6-α4), B (β9-α7) and C (the C-terminal region) are the most important because they are related to cofactor binding and/or substrate specificity (Jez et al., 1997). Figure 3 shows a sequence alignment including five representatives of L-GDH from different species of plant together with seven AKRs from different families which have known 3D structures available. L-GDHs have a short Loop-A, similar to that seen in the AKR3, AKR5, AKR7 and AKR11 families, but different when compared to AKR1, AKR2, and AKR4. AKRs1-5 have a short Loop-B, L-GDHs a medium-sized Loop-B and AKR7/AKR11 a long Loop-B. Loop-C is very variable among all the sequences. Like most AKRs, L-GDHs have two α-helices external to the barrel, the H1 helix within Loop-B and the H2 helix between α8 and Loop-C (both helices are conserved in most AKRs). The lack of a description of the 3D structure for L-GDH up until now has precluded its adequate classification within the AKR superfamily.

**Figure 2.**
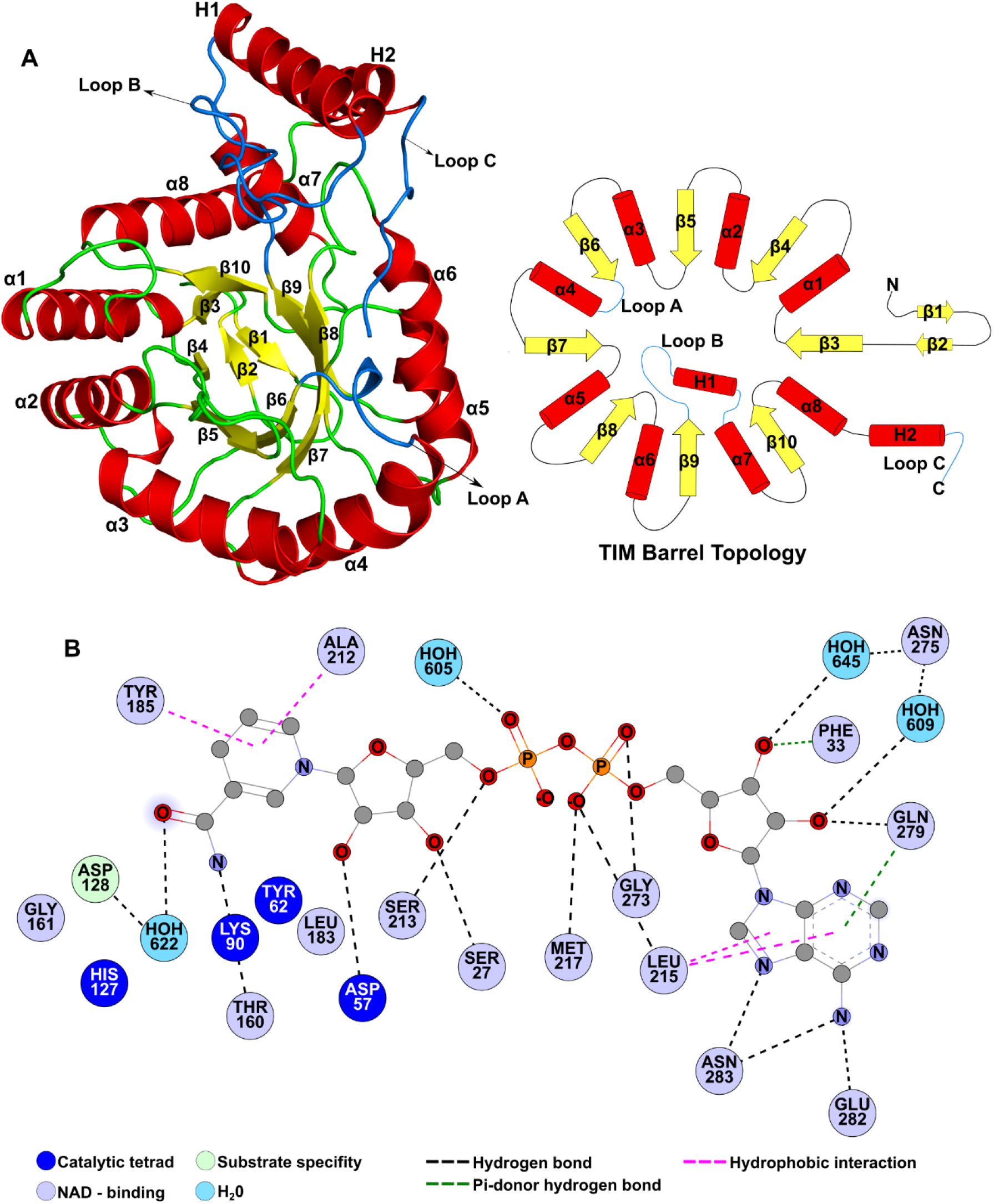
*So*GDH fold and NAD^+^ binding interactions. (A) Cartoon representation (left) and topology diagram (right) for the structure of *So*GDH which shows a (β/α)_8_--barrel (TIM-barrel) fold, characteristic of the AKR superfamily. This fold has 8 β-strands (yellow) interspersed by 8 α-helices (red). At the bottom of the barrel, at the N-terminus of the polypeptide chain, there is a β-hairpin composed of strands β1 and β2. In addition, *So*GDH presents two helices (H1 and H2) external to the barrel, which together with the loops Loop-A, Loop-B and Loop-C are important for the classification of the members of AKRs. (B) Shows the network of interactions with the NAD^+^ cofactor. Specific features are highlighted according to the following colour code; in dark blue the amino acids of the catalytic tetrad; in light blue positions conserved in all AKRs which interaction with NAD^+^; in light green amino acids important for determining substrate specificity and in cyan, water molecules. Dashed lines indicate hydrogen bonds (black), hydrophobic interactions (purple), and pi-donor hydrogen bonds (dark green). Figures were generated with PyMol v2.05 (Schrödinger, LLC) and Discovery Studio Visualizer V21.1.0 (BIOVIA, Dassault Systèmes).

**Figure 3.**
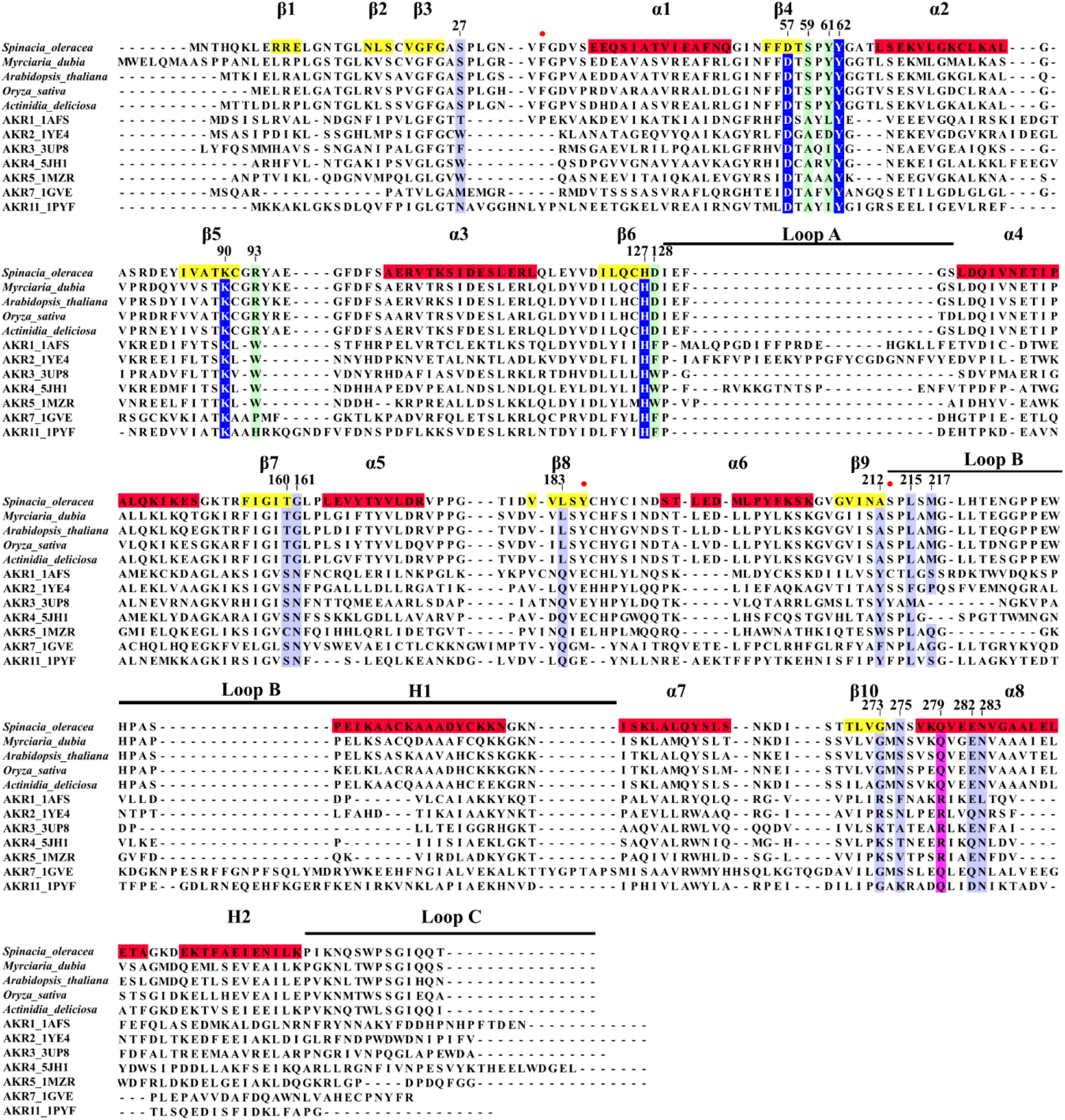
Multiple sequence alignment between L-GDHs and AKRs. Five L-GDH sequences from different species were aligned with seven representatives of different families of AKRs. Using the spinach *So*GDH sequence, the position of secondary structure elements (strands in yellow and helices in red) are indicated. Likewise, the numbering adopted for the description of the amino acid positions was based on *So*GDH. In dark blue the amino acids of the catalytic tetrad are highlighted, in light blue those responsible for binding to NAD^+^, in light green the amino acids important in determining substrate specificity and in purple a conserved position in ARKs that confers a preference for the cofactor (NAD^+^ or NADP^+^). Positions with a red ball represent residues that interact with NAD^+^ which are only observed in the L-GDH structures presented here.

### NAD^+^ binding

AKRs are characterized by binding pyridine nucleotide coenzymes, commonly NAD(P). Likewise, they present a conserved oxidation-reduction mechanism using a catalytic tetrad composed of highly conserved amino acids (Asp, Lys, Tyr, and His), having specificity for sugar or steroid substrates (Jez et al., 1997). It is well established that L-GDH has a specificity for NAD^+^ as its cofactor (rather than the more common NADP^+^) and that the sugar L-gal is its preferred substrate (Gatzek et al., 2002). In order to fully explore the differences between L-GDH and other members of the AKR protein superfamily, it was also necessary to solve the structure of *So*GDH in the form of its complex with NAD^+^. This structure will be referred to here as *So*GDH-NAD^+^.

The crystal form of *So*GDH-NAD^+^ had two molecules in the asymmetric unit. However, PISA analyses (Krissinel and Henrick, 2007) indicate that this apparent homodimer is not expected to exist under physiological conditions and is merely the result of crystal packing. This is consistent with the SEC-MALS results presented previously. Indeed, to the present date, the only members of the AKR superfamily which are known to be homodimers are aflatoxin dialdehyde reductase (AKR7A1), NAD(P)H dependent xylose reductase (AKR2B5), and the tyrosine auxotrophy suppressor protein (Kavanagh et al., 2003; Kozma et al., 2002; Obmolova et al., 2003). A similar dimerization interface using helices α5, α6, H2, and Loop-C is observed in all of these structures (figure S3). On the other hand, in *So*GDH-NAD^+^ the contact interface between the two monomers of the asymmetric unit is completely different and involves the loop β5-α3, Loop-B, and Loop-C of chain A and the helices α5, α6 and loop α5-β8 of chain B (figure S3). Consequently, the two molecules in the asymmetric unit of *So*GDH-NAD^+^ interact merely as the product of crystal contacts and has no physiological relevance. This observation is reinforced by the apo structure of *So*GDH which crystallizes with a monomer in the asymmetric unit in a space group which lacks 2-fold rotation axes and therefore dimers are impossible. The sum total of the evidence presented here indicates that spinach *So*GDH is a monomeric protein (at least under the conditions used in the present work) and not a homodimer as reported by Mieda et al (Mieda et al., 2004).

*So*GDH has the conserved catalytic tetrad observed in AKRs: Asp57, Tyr62, Lys90, and His127 (figure 4). The tetrad is also part of a group of residues that interact directly with NAD^+^ (figure 2B). This strongly suggests that L-GDH follows the same basic catalytic mechanism described for other AKRs but with a preference for favouring the oxidation (dehydrogenation) reaction rather than reduction (see below). This leads to the production of L-galactono-1,4-lactone from L-galactose with the concomitant reduction of NAD^+^ to NADH.

**Figure 4.**
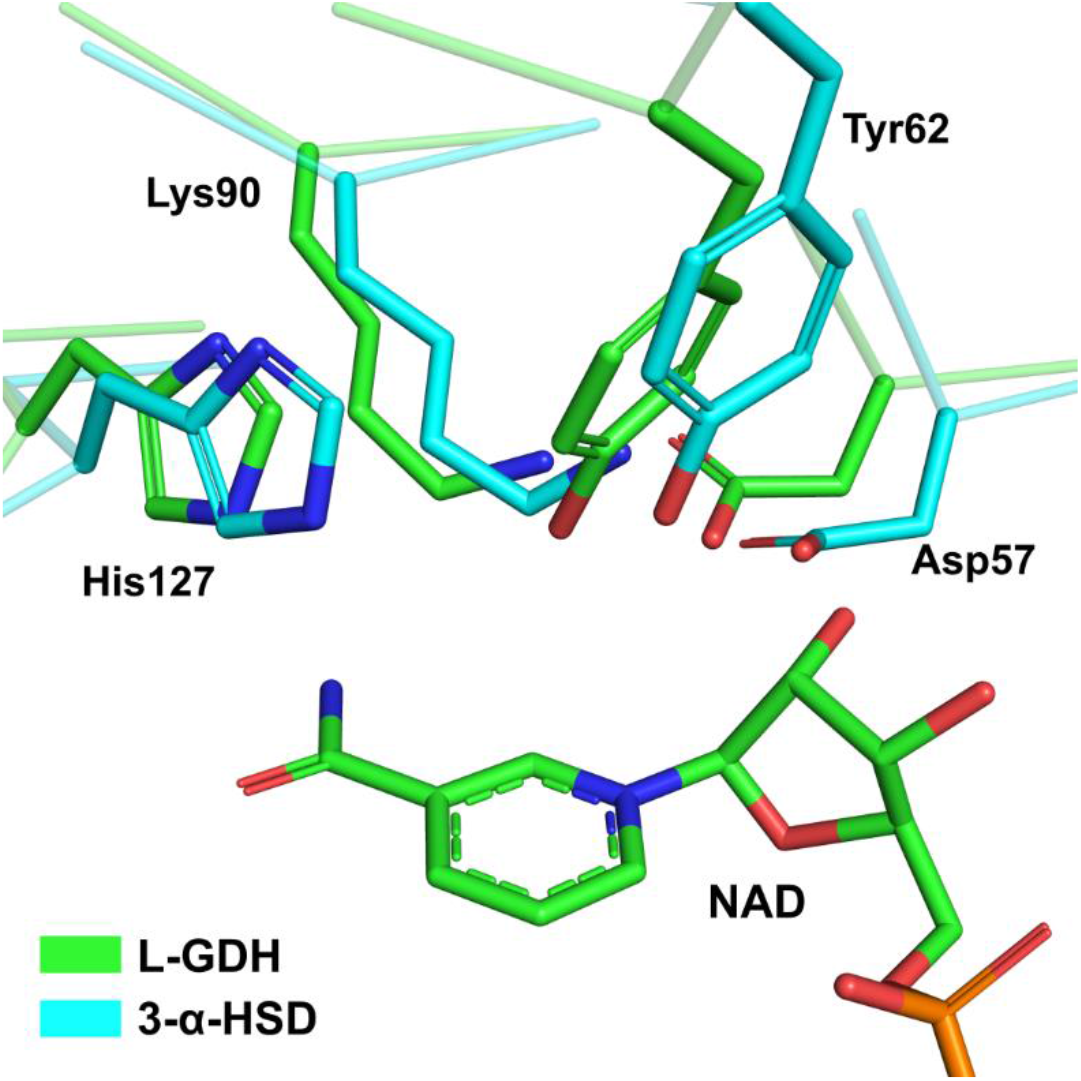
The active site in *So*GDH. A superposition between the active site of *So*GDH (green) and 3-α-hydroxysteroid dehydrogenase (a representative of the AKR1 family, in cyan) shows the catalytic tetrad (Asp27, Tyr52, Lys90 and His127) conserved in L-GDH. 3-α-HSD PDB code: 1AFS.

AKRs in general have a group of conserved residues responsible for NAD(P) binding, the most conserved being Asp57, Ser160, Asn161, and Gln183 (figure 3) (Bennett et al., 1997; Jez et al., 1997). Of these residues, *So*GDH retains only Asp57, while at position 160 it has a Thr, at 161 a Gly and at 183 a Leu. These changes lead to the loss of two hydrogen-bonds with the nicotinamide ring, forming new Van-der Waals interactions with Gly161 and Leu183 (figure 5A e 5B). The substitution of the aromatic residue at position 212 by Ala in *So*GDH also stands out (figure 3). These aromatic amino acids, in AKRs occupy a position on the underside of the nicotinamide ring as seen in figure 5A and are a notable feature of the cofactor binding site (Bennett et al., 1997; Jez et al., 1997). Given its structural and functional importance, the lack of this aromatic residue in L-GDH is, at first glance, surprising. However, in L-GDHs, the pi-stack interaction is recovered with Tyr185, a conserved amino acid in these enzymes (Figure 5B and figure 3). This appears to be a classical example of a compensating substitution. A hydrogen bond between Thr27 and the ribose ring on the nicotinamide side of the cofactor in AKRs is maintained via Ser27. Additionally, in *So*GDH, a new hydrogen bond is observed between Ser213 and the phosphate group on the nicotinamide side of the cofactor (PN) (Figure 5B).

**Figure 5.**
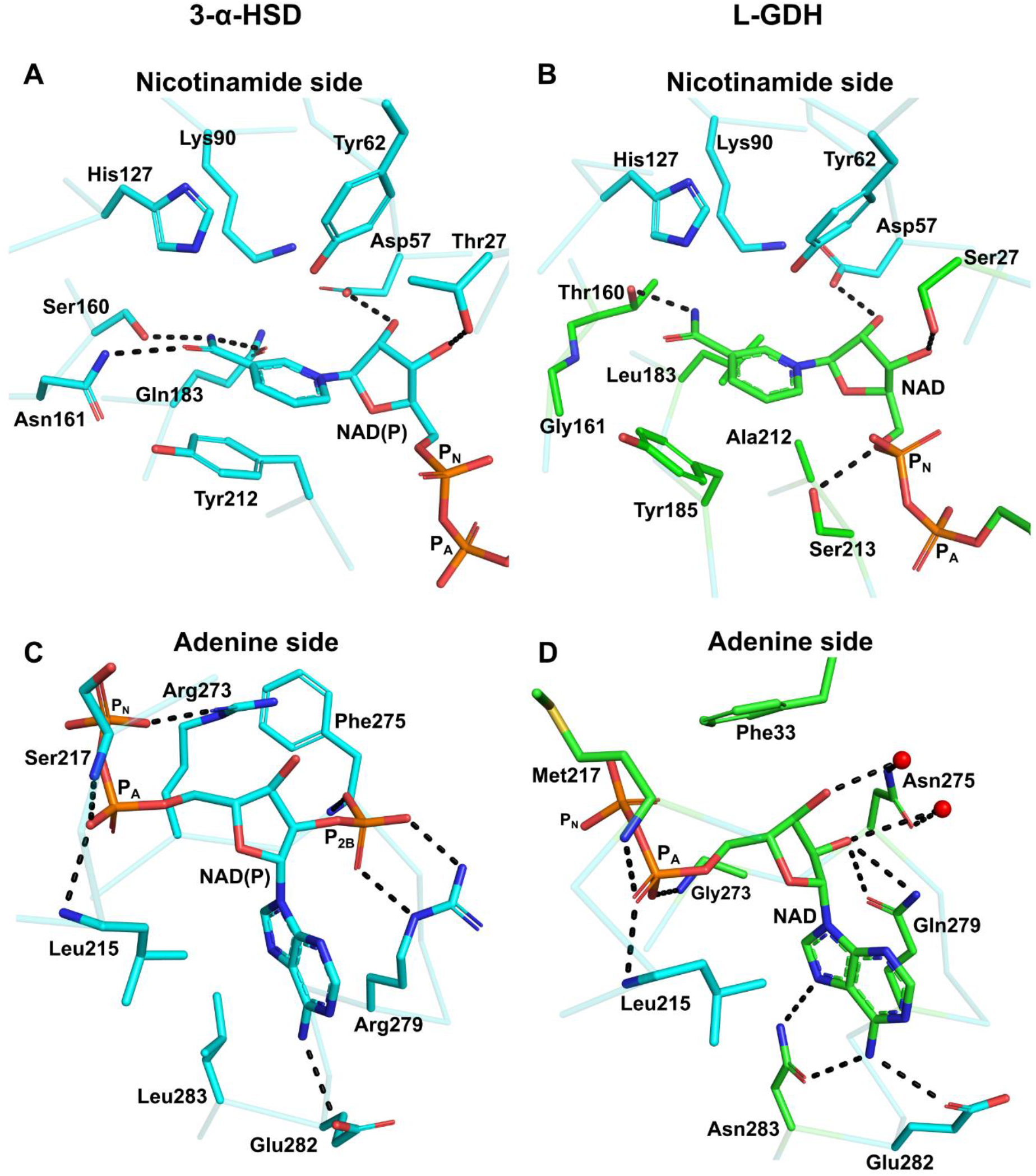
Comparison between the NADP^+^ binding interactions in AKRs and NAD^+^ binding in *So*GDH. For a better understanding, the interactions have been divided into the nicotinamide and adenine sides which are treated separately. (A) and (C) show the interactions made with NADP^+^ in 3-α-HSD (PDB identifier: 1AFS). On the nicotinamide side (A), three hydrogen bonds are observed between Ser160, Asn161 and Gln183, which together with the pi-stacking formed by Tyr212, correctly orient the nicotinamide ring of NADP^+^ for catalysis. On the adenine side (C), the interaction between Arg279 (conserved in AKRs) and the P2B phosphate stands out and confers specificity for NADP^+^. On the nicotinamide side of *So*GDH (B) the hydrogen bonds are lost, as well as the pi-stack at position 212. However, the orientation of the nicotinamide ring is maintained by the hydrophobic interaction of Leu183 and the pi-stacking of Tyr18. In addition, a hydrogen bond is maintained between Thr160 and the nicotinamide ring. On the adenine side (D), we observe that the absence of P2B allows for interaction via water molecules between Asn275 and the ribose. Likewise, Gln279 interacts directly with the ribose. A hydrophobic interaction involvingPhe33 and two new hydrogen bonds between Asn283 and the adenine appear in *So*GDH. PN: phosphate on nicotinamide side of the cofactor, PA: phosphate of adenine side. On comparing (B) with (A) or (D) with (C), conserved residues have retained the colour blue whereas substitutions are shown in green.

Changes in the interaction between the enzyme and the cofactor are also evident on the adenine side of the NAD^+^. Lys/Arg273 and Arg279 confer NADP^+^ specificity to the majority of AKRs (figure 5C) (Jez et al., 1997). In *So*GDH, position 273 is replaced by a Gly and 279 by a Gln (figure 5D). In AKRs, Lys/Arg273 forms a hydrogen bond with PN (the phosphate on the nicotinamide side of the cofactor). However, in *So*GDH, Gly273 makes a hydrogen bond using its main chain to the phosphate on the adenine side (PA). However, most important appears to be the loss of Arg279 in GDHs (figure 3). Arg279 forms hydrogen bonds with the P2B phosphate moiety on the adenine side ribose, a specific interaction for enzymes which employ NADP^+^ as the cofactor (figure 5C) (Jez et al., 1997). In *So*GDH, the lack of Arg279 means that the enzyme is unable to compensate the formal charge on P2B rendering L-GDHs specific for NAD^+^ rather than NADP^+^. Nevertheless, the substitution by Gln still allows for direct interaction with one of the ribose hydroxyl groups on the adenine side (figure 5D).

In AKRs, residue 275 is highly variable (figure 3) but this does not influence its interaction with P2B because this occurs via its main chain amine (Jez et al., 1997) (figure 5C). However, in GDHs, the homologous residue, Asn275, is highly conserved, and due to the absence of P2B, Asn275 can interact with the ribose moiety via two water molecules (figure 5D). In the same region of *So*GDH, a new interaction is observed. Phe33 makes a hydrophobic contact and a pi-donor hydrogen bond with the ribose on the adenine side (figure 5D). Furthermore, Leu283, which makes hydrophobic contact with the adenine in AKRs, is substituted by Asn in *So*GDH. This allows for the formation of two additional hydrogen bonds with the adenine ring, further stabilizing the cofactor (figure 5D).

Overall, as figure 5 makes clear, the nicotinamide side of the cofactor binding site is much more conserved than the adenine side. In the case of the former, this is not unexpected given the need, in both cases, for the binding site cavity to maintain the orientation of the nicotinamide moiety orientated for productive catalysis. However, in the case of the latter, although some aspects of the binding site are preserved when comparing AKRs and L-GDHs, overall the cavity has been radically altered in order to accommodate NAD^+^ rather than NADP^+^. The preference for NAD^+^ as the cofactor for L-GDH is probably related to the direction of the reaction being catalyzed which, in order to generate AsA as the final product, must be in the oxidative direction. As pointed out by Barski et al (2008) this is different to most AKRs which generally favour reduction. These enzymes use NADPH as the cofactor which is more abundant in the cytoplasm than the oxidized form NADP^+^, thereby favouring the reduction reaction (Barski et al., 2008). For NADH/NAD^+^ the reverse is true with NAD^+^ being the predominant species. By preferring NAD^+^ as the cofactor, L-GDH favours the oxidative production of L-galactono-1,4-lactone from L-galactose and thereby the effective production of ascorbic acid.

### Substrate specificity

Thus far, we have been unsuccessful in obtaining crystals of *So*GDH bound to the natural substrate (L-Gal), a substrate analogue or an inhibitor. Nevertheless, comparisons with crystal structures of AKRs complexed to inhibitors are still possible in order to investigate the structural basis for differences in substrate specificity. For this purpose, we will use as an example of AKRs 3-α-hydroxysteroid dehydrogenase (3-α-HSD) and human aldose reductase (ADR), both bound to inhibitors (figure 6). AKRs present amino acids within the substrate binding pocket which are responsible for differentiating between a wide range of steroids and sugar substrates (light green in figure 3) (Bennett et al., 1997; Jez et al., 1997). Figure 6 shows that in the active site of the three enzymes, Asp57, Lys90, Tyr62, and His127 (the catalytic tetrad) are fully conserved (figure 6). Trp228 from Loop-B is essential for stability rather than substrate discrimination and is also conserved in all three proteins (Bennett et al., 1997; Jez et al., 1997). Furthermore, Ala59 and Trp93 play the same role in substrate stabilization in both 3-α-HSD and ADR (Figure 6A and 6B). However, in *So*GDH these positions are occupied by Ser59 and Arg93, which could be important for substrate discrimination since they have the potential to form new hydrogen bonds with the hydroxyl groups on the L-Gal substrate (figure 6C). Amino acids at positions 61 and 128 are involved in the discrimination between sugars and steroids (Jez et al., 1997). At position 61, the hydroxysteroid dehydrogenases (HSDs) have Leu or Ile, while aldose reductases (ADRs) have Val. Differences in this position can alter the topology of the binding site to accommodate substrates of different sizes. Furthermore, at position 128, HSDs have Phe while ADRs have a conserved Trp. The presence of the indole nitrogen allows discrimination between the sugar and the steroid, controlling the accessibility of the substrate or its orientation in the active site (figure 6A and 6B) (Jez et al., 1997). Spinach *So*GDH exhibits what would appear to be highly significant differences at these two positions, which are occupied by Tyr61 and Asp128 (figure 6C). Tyr61 could potentially perform a stacking interaction with the sugar ring or hydrogen bonds with the substrate via its hydroxyl group. Asp128, on the other hand, has the potential to form a pair of hydrogen bonds with adjacent hydroxyls on L-Gal aiding in establishing the correct orientation of the substrate whilst discriminating between different sugars. Furthermore, galactose-binding enzymes often use an aromatic amino acid (Trp, Phe, or Tyr) to correctly orient the galactose substrate within the active site (Sujatha et al., 2004). GDHs, including *So*GDH have a conserved Tyr185 that differentiates them from the remaining AKRs (figure 3). In addition to restoring the interaction with the NAD^+^ nicotinamide ring, this tyrosine could also play a role in establishing the specificity of the enzyme for L-galactose and orienting it correctly in the binding pocket (figure 6C).

**Figure 6.**
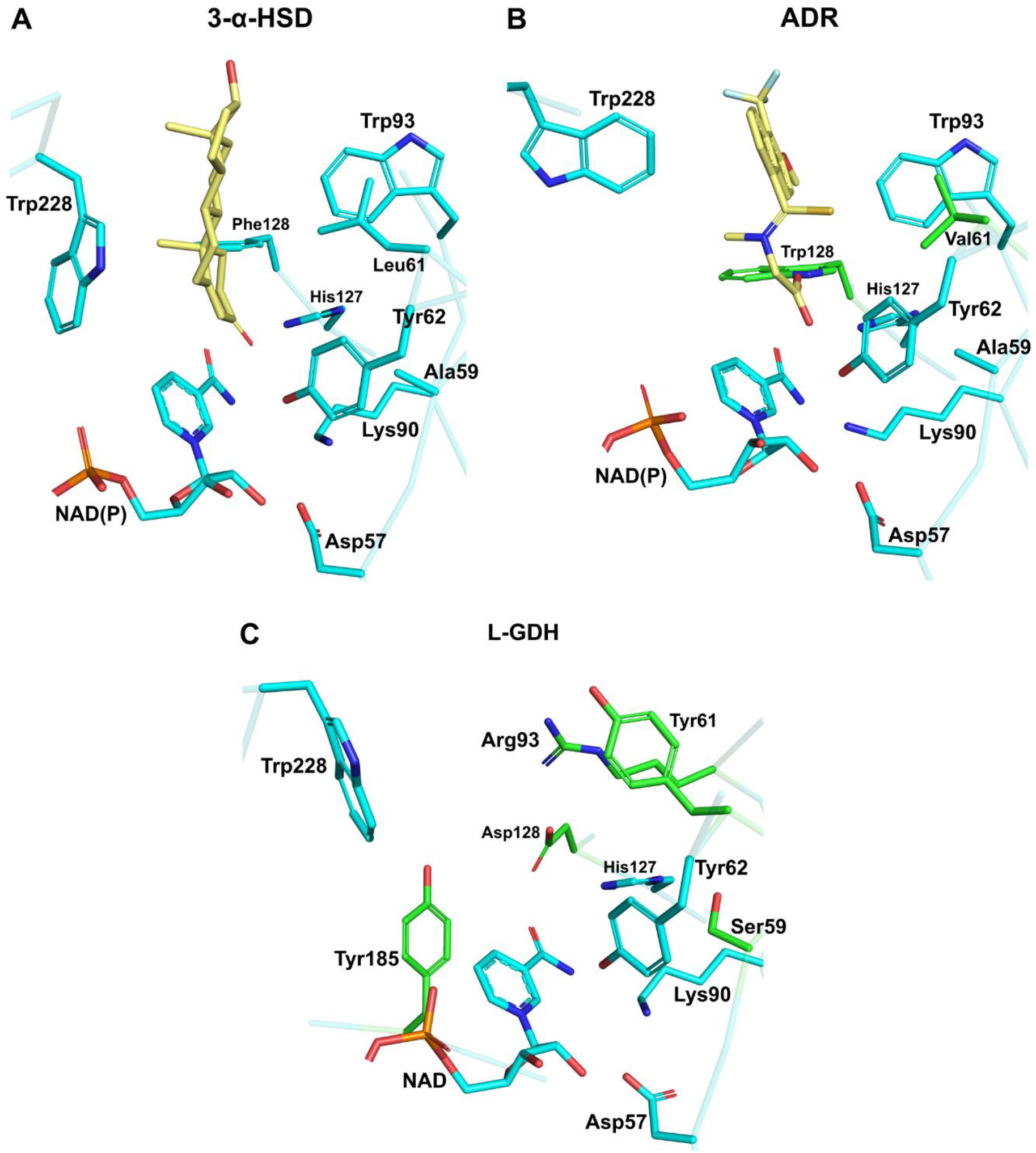
Substrate specificity in AKRs and SoGDH. (A) 3-α-HSD bound to testosterone (yellow), showing amino acids important for substrate binding. Leu61 and Phe128 are specific for AKRs that bind steroids. (B) Human aldose reductase (ADR) bound to the inhibitor tolrestat. Val61 and Trp128 (green) are specific for AKRs that bind sugars. (C) The empty active site as observed in the *So*GDH-NAD^+^ complex. Despite *So*GDH having a sugar as its natural substrate, it presents a Tyr at position 61 and an Asp at 128. It also has differences in positions 59 (Ser) and 93 (Arg). Tyr185 appears to be specific for L-GDH, since galactose-binding proteins use an aromatic amino acid to orient the ligand to the active site. 3-α-HSD PDB code: 1AFS, ADR PDB code: 2FZD.

Loop-C is essential for determining substrate specificity in ARKs (Jez et al., 1997). However, in both forms of *So*GDH reported here, loop-C is not oriented towards the active site due to the absence of substrate and we are so far unable to shed light on its role in the case of GDHs. In summary, despite the absence of a complex with the substrate or a substrate analogue, the structures presented here provide the basis for testing, by site directed mutagenesis, those residues which are most likely to be involved in substrate recognition and selectivity.

Despite the conservation of the (β/α)_8_ fold, the significant differences observed at both the cofactor and substrate binding sites would appear to justify the classification of GDHs as a new family within the AKR superfamily. This is borne out by a phylogenetic analysis of representative sequences of the existing families taken from the University of Pennsylvania PKR database (https://hosting.med.upenn.edu/akr/) which shows GDHs to cluster into a monophyletic group (figure S4).

### Open and closed state

AKRs have an NAD(P)-dependent opening and closing mechanism, which allows for the formation of a tunnel that maintains NAD(P) bound. This transitory structure is formed after cofactor binding by a conformational change in Loop-B and Loop β3-α1 (Bennett et al., 1997). The interactions that allow tunnel formation are different in HSDs and ADRs. ADRs differ by the presence of a salt bridge involving Asp227, Lys35, and Lys273 and hydrophobic interactions made by Trp27 (Loop-β3-α1) and Pro226-Trp228 (Loop-B). Due to the mutation of Asp227 to a Ser, HSDs lack the salt bridge that helps stabilize the closed state. Consequently, in HSDs, the tunnel is stabilized by different interactions formed between several residues from Loop-β3-α1 (Pro33, Glu34, Lys35) and Loop-B (Asp225, Lys226, Thr227, and Trp228) (Bennett et al., 1997).

Spinach *So*GDH presents both the open and closed states as a function of NAD^+^ binding. An overlay of the two structures yields an RMSD of 0.66 Å for 318 aligned Cα atoms with the regions showing the greatest divergence being Loop-β3-α1 and Loop-B (important for tunnel formation) (figure 7). Despite *So*GDH having a sugar substrate, like ADRs, they do not have a salt bridge in the tunnel because at position 273, a Gly substitutes for the Lys/Arg conserved in ARKs. Tunnel formation and stabilization are due to hydrophobic interactions between Val32-Phe33 (Loop-β3-α1) and Pro226-Trp228 (Loop-B), as well as a hydrogen bond between the indole nitrogen of Trp228 and the Val32 main chain (figure 7C). Although there is significant variation amongst AKRs and GDH in the details of the interactions responsible for tunnel formation, this appears to be a common mechanism for all such enzymes indicating that NAD^+^ binding induces a conformational change to the enzyme which is presumably essential for competent catalysis.

**Figure 7.**
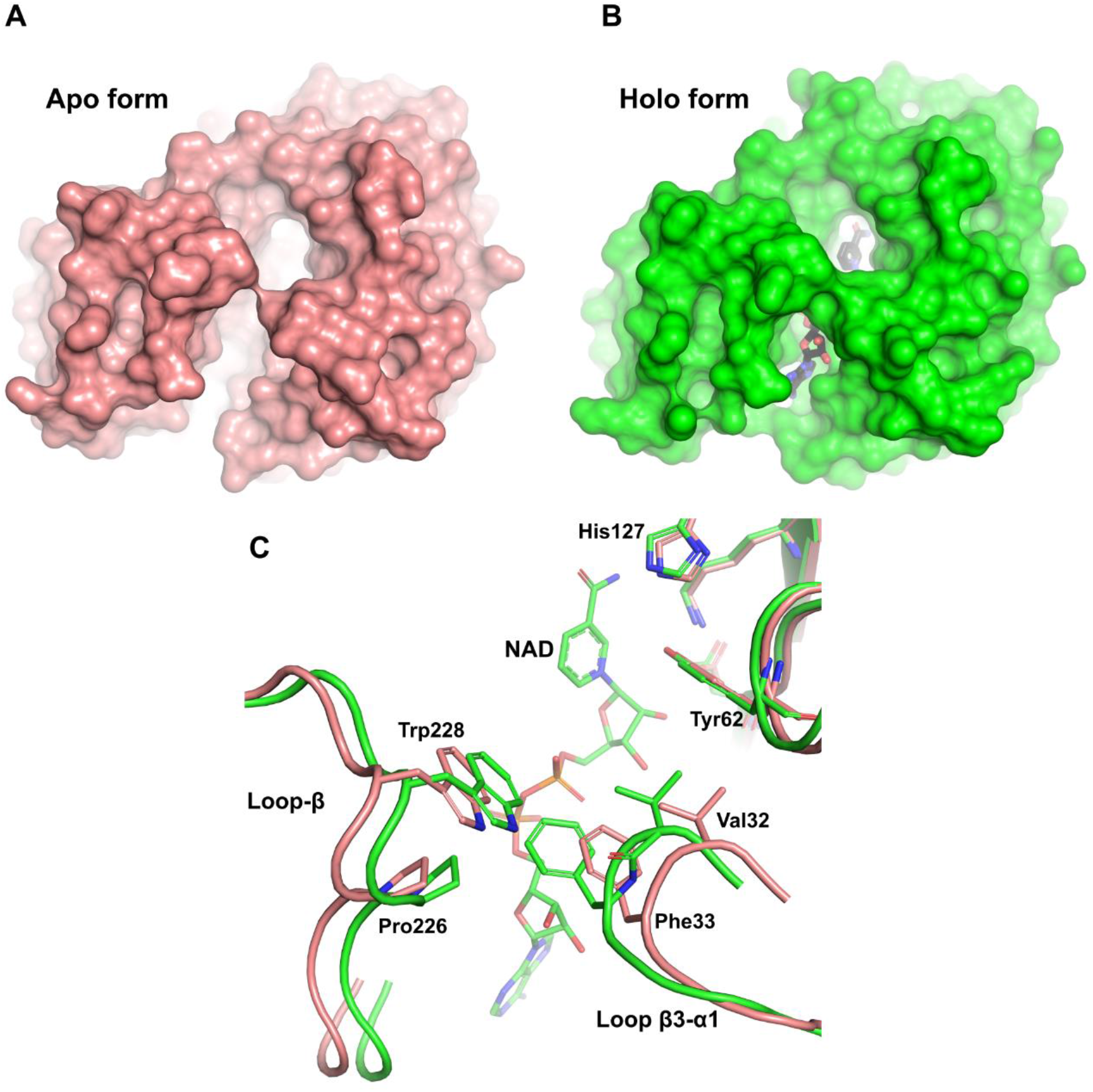
The open and close states of *So*GDH. (A) The apo form of *So*GDH (salmon) shows the open state, with little contact between Loop-B and Loop-β3-α1. (B) The holo form of *So*GDH-NAD^+^ (green) shows the closed state, forming a tunnel generated by the contact made between Loop-B and Loop-β3-α1 (NAD^+^ is shown in black within the tunnel at the bottom). (C) A superposition between *So*GDH apo and holo forms shows a close-up of the Loop-B and Loop-β3-α1 regions. NAD^+^ binding results in hydrophobic contacts between Val32, Phe33, Pro226, and Trp228, as well as hydrogen bonding between the indole nitrogen of Trp228 and the Val32 main chain.

### L-GDH structure comparison

The primary structures of L-GDHs show high levels of sequence identity among different plant species (percentage identity of 76%), preserving key amino acids for NAD^+^ binding and those deduced from the discussion above to be important for L-galactose substrate specificity (figure 3). Therefore, they would be expected to have a well-conserved three-dimensional structure, independent of species.

The RCSB protein data bank currently only has available the three-dimensional structures of rice *Os*GDH in its apo (7EZI) and holo (7EZL) states. However, there is no published description of these structures. As anticipated, rice *Os*GDH has the highly conserved (β/α)_8_-barrel fold similar to spinach *So*GDH. The apo form of both proteins has an RMSD of 0.45 Å for Cα atoms and the holo-structures 0.46 Å. Significant differences only exist in the Loop-β3-α1 and Loop-β4-α2 regions (figure 8A). The importance of the Loop-β3-α1 region for tunnel formation and the fact that the amino acid sequences in these regions is almost identical in both enzymes makes these differences to be of interest and worthy of further investigation. At first glance, such similar sequences would not be expected to adopt such radically different conformations.

**Figure 8.**
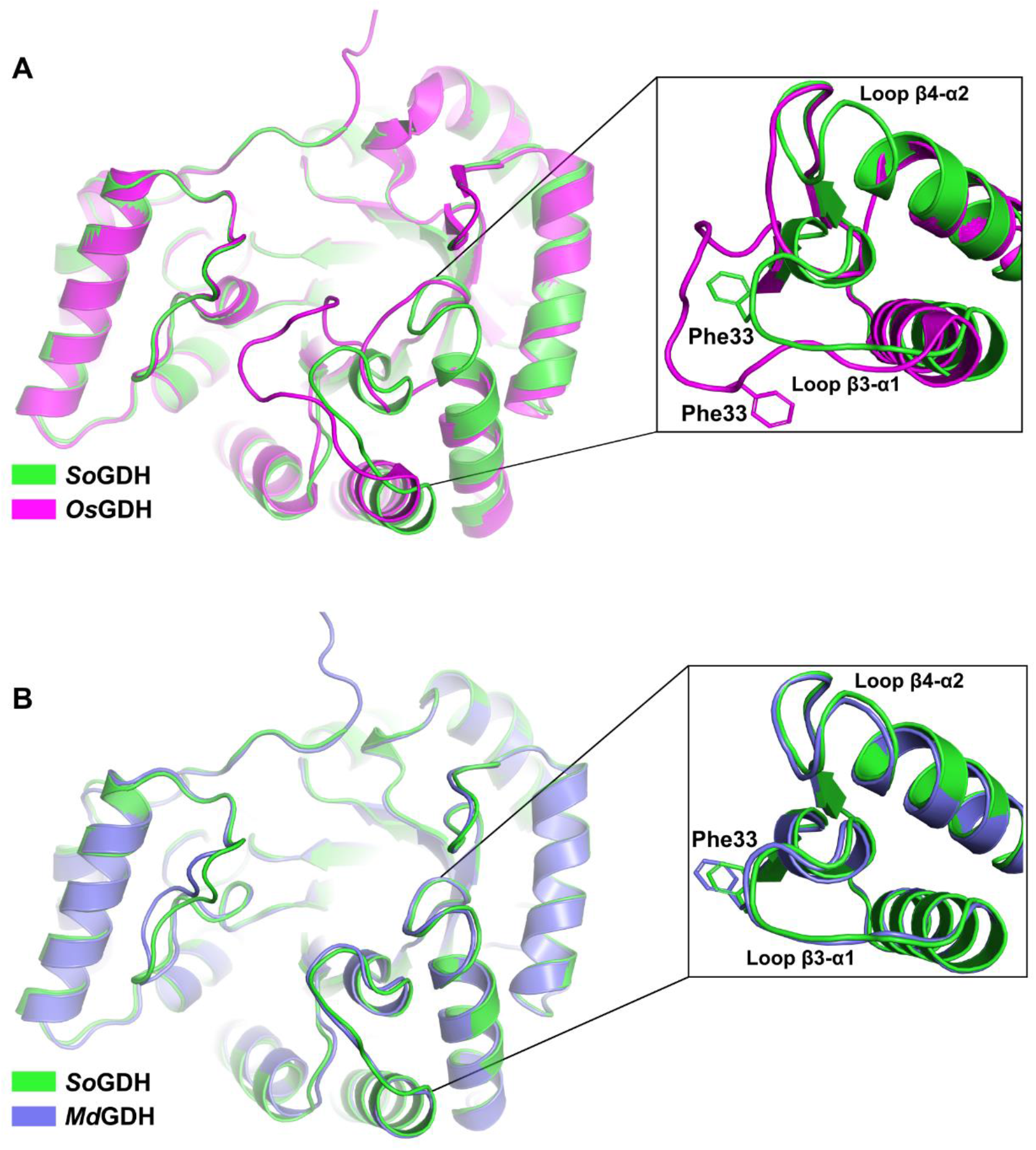
L-GDH structure comparisons. (A) A superposition between spinach *So*GDH (green) and rice *Os*GDH (purple) crystal structures in the apo form. Despite having an RMSD of 0.45 Å for Cα atoms, the orientation of Loop-β3-α1 and Loop-β4-α2 are very different (with a displacement of approximately 6Å). (B) Superposition between the crystal structure of spinach *So*GDH (green) and the Alphafold2 model for camu-camu *Md*GDH (light blue). Loop-β3-α1 and Loop-β4-α2 show the same orientation. Rice *Os*GDH PDB code: 7EZI.

The different conformation of Loop-β3-α1 in the rice enzyme leaves Phe33, necessary for tunnel formation, in an orientation opposite to that observed in spinach *So*GDH (figure 8A, inset). In addition, the shift in position of Loop-β3-α1 (∼6 Å) towards Loop-B induces the formation of a tunnel even in the apo form, something not normally observed in AKRs or in *So*GDH (figure 8A). The quality of the electron density map in Loop-β3-α1 and Loop-β4-α2 of spinach *So*GDH in both the apo and holo forms leaves no room for ambiguity and shows that the conformation reported for the Loop-β3-α1 region has been correctly interpreted (Figure S5). However, analyzing the electron density map of *Os*GDH it becomes evident that this region could have been misinterpreted. Furthermore, it appears to have a knock-on consequence for the neighbouring Loop β4-α2 which also adopts a conformation radically different to that seen in *So*GDH.

Since we have thus far been unsuccessful in obtaining diffraction quality crystals of GDH from camu-camu (*Md*GDH) we performed its structure prediction using Alphafold2 (Jumper et al., 2021). The resulting model for camu-camu *Md*GDH presents a very similar structure to that of *So*GDH with an RMSD of 0.56 Å for Cα atoms. In *Md*GDH, an open state is observed with Loop-β3-α1 and Loop-β4-α2 in the same positions as observed in spinach *So*GDH (figure 8B), and very different to that seen in the structure for rice *Os*GDH. There are no notable differences between *So*GDH and *Md*GDH in the vicinity of the substrate and NAD^+^ binding sites implying that it would be expected that they would present similar kinetic properties as well observe experimentally (Table 1).

Notwithstanding the fact that AlphaFold2 predictions are still under scrutiny by the scientific community, our results strongly suggests that the conformation described here for the loops in apo SoGDH is correct and would be expected to be conserved in other GDHs. Since this region is involved in tunnel formation and therefore is of functional significance, it appears that the structural interpretation of this region in apo *Os*GDH is worthy of revisiting.

## Conclusion

In this work we contribute to closing the knowledge gap about functional and structural aspects of critical enzymes involved in vitamin C biosynthesis in plants, particularly of the D-mannose/L-galactose pathway. In this context, we report enzyme kinetic parameters (e.g., *kcat*, catalytic efficiency, etc.) for L-GDH from two different plant species and describe it crystal structure for the first time in both its apo and holo forms. This reveals the structural basis for the preference for NAD^+^ as the cofactor (important for driving the reaction in the direction of ascorbic acid synthesis) and that the enzyme undergoes a similar conformational change on cofactor binding to that observed in AKRs in general. This leads to the burial of NAD^+^ inside a binding site tunnel with the nicotinamide directed towards the region of the catalytic tetrad. Despite the absence of a crystal structure of the enzyme with L-Gal it has been possible to identify residues which are likely involved in imbuing selectivity for its natural substrate. These include Tyr61, Tyr185, Ser59 and Asp128 and their identification opens up new perspectives for future experiments.

The crystal structure of spinach *So*GDH shows its great similarity with aldo-keto reductases which is based in the classical (β/α)_8_-barrel fold. This reveals a common evolutionary origin despite the preference for most such enzymes to favour reductive catalysis rather than oxidation, as is the case for GDHs. Hence the use of NAD^+^ as the cofactor rather than NADPH. The identification of the residues which favour both NAD^+^ and L-Gal binding suggests that L-GDH could be considered to form a new group within the AKR superfamily.

The regulatory mechanism behind the control of AsA synthesis in plants is an important open question. According to our results, both *Md*GDH and *So*GDH are refractory to inhibition by AsA, indicating that in camu-camu and spinach plants, the catalytic activity of L-GDH is not regulated by feedback inhibition. Previous reports to the contrary may have been artifacts introduced by a change in pH rather than competitive inhibition. These results highlight the potential biotechnological applications for the use of GDH in the production of vitamin C and the need for further investigation into the *in vivo* control of vitamin C production, a complex issue given the number of different synthetic routes employed in many plants.

## Materials and Methods

### Plasmid construction

The CDS for L-galactose dehydrogenase from *Myrciaria dubia* (*Md*GDH) (GenBank accession No. OK632632.1) was optimized using the GeneOptimizer software suite (Raab *et al*. 2010), then synthetized by GeneArt® (Invitrogen, Carlsbad, CA, USA) and ligated into the plasmid expression vector pET151/D-TOPO. The synthetic gene for L-galactose dehydrogenase from spinach (*Spinacia oleracea, So*GDH) (GenBank accession No. AB160990.1) was ligated into the plasmid expression vector pET-28a, which was purchased from FastBio (São Paulo, Brazil).

### Protein expression and purification

*Escherichia coli* BL21 Rosetta™ (DE3) cells harboring the constructs were grown at 37 °C in LB medium supplemented with ampicillin (50 μg·ml^−1^) and chloramphenicol (34 μg·ml^−1^) for *Md*GDH, and kanamycin (30 μg·ml^−1^) and chloramphenicol (34μg·ml^−1^) for *So*GDH. Once the OD600nm = 0.5 – 0.6 was reached, the culture was cooled to 20 °C and protein expression induced by addition of 0.3 mM isopropyl 1-thio-β-D-galactopyranoside (IPTG). After 16 h, cells were harvested by centrifugation at 10,000 g for 45 min at 4 °C and suspended in lysis buffer (150 mM NaCl, 20 mM Tris-HCl pH 7.5). The soluble fraction, after cell lysis by sonication, was isolated by centrifugation at 16,000 g for 45 min at 4 °C. This was then loaded onto a column with 5 ml His60 Ni-Superflow Resin (Clontech) previously equilibrated in the lysis buffer. Subsequently, the resin was washed with 5 column volumes of lysis buffer and then again with 5 column volumes of lysis buffer containing 50 mM imidazole. Bound proteins were eluted from the resin using 2 column volumes of lysis buffer containing 250 mM imidazole. The next purification step, size exclusion chromatography, was performed using a Superdex 200 XK16 column (GE Healthcare) pre-equilibrated in lysis buffer. The purity of the eluted enzymes recovered from both purification steps was evaluated by SDS-PAGE. The desired concentration of proteins was achieved by centrifugation at 800 g, using an Amicon® Ultra (molecular weight cut-off 30 kDa) centrifugal filter device (Merck Millipore, Darmstadt, Germany). Samples were kept frozen at -80 °C for future use.

### Size exclusion chromatography coupled with multi-angle light scattering

The oligomeric state of *Md*GDH and *So*GDH was evaluated by SEC-MALS using a three-angle light scattering detector miniDAWN® TREOS® (Wyatt Technology, Santa Barbara, CA, USA) and Optilab T-rEX differential refractometer (Wyatt Technology, Santa Barbara, CA, USA). For the size exclusion chromatography, this system was coupled to an HPLC (Waters, Milford, MA, USA) consisting of a pump and controller (Waters 600, Milford, MA, USA). 50 μl of each sample (at a concentration of 3 mg·ml^−1^) were loaded onto either a Superdex 75 or 200 HR 10/300 GL columns (GE Healthcare, Madison, WI, USA), equilibrated in 25 mM Hepes (pH 7.8), 300 mM NaCl and 5 mM MgCl2. The data collection and analysis were performed via Wyatt ASTRA 7 software (Wyatt Technology Corporation, Santa Barbara, CA, USA).

### Enzyme assays

Enzyme kinetics, inhibition assays and the influence of pH were performed in a SpectraMax Plus 384 Microplate Spectrophotometer (Molecular Devices, Sunnyvale, California, USA). The conversion of NAD^+^ to NADH (ε = 2650 M^−1^cm^−1^) was monitored spectrophotometrically, in a 96 well UV-transparent microplate, by recording the absorbance at 324 nm every 6 s for 5 min at 26 °C. Initial enzyme velocities were estimated as the maximum linear rates of absorbance increase at 340 nm. mAU values were converted to molar concentration using the Beer-Lambert equation A = ε x 𝓁x c, where: A is the absorbance, ε is the molar extinction coefficient of NADH at 324 nm, 𝓁is the optical path length in cm and c is the molar concentration of NADH. Then, these values were converted to reaction rate units (concentration/time).

### Enzyme Optimal pH determination

The optimal pH for *So*GDH and *Md*GDH was determined in a pH range from 4 to 11. Catalytic activity was measured as mentioned above, in 200 μL reaction mixture in a universal buffer (a mixture of three tri-protic acids: citric acid, boric acid, and phosphoric acid) (Ganesh et al., 2017) containing 200 μM of NAD^+^, L-galactose (L-gal, 1 mM for *So*GDH and 2 mM for *Md*GDH) and the purified recombinant enzyme was added to a final concentration of 100 nM.

### Enzyme Kinetics analysis

Recombinant *So*GDH and *Md*GDH were assayed in a 200 μL reaction mixture containing 100 mM Tris-HCl pH 7.0, 200 μM of NAD^+^, L-gal (1 mM for *So*GDH and 2 mM for *Md*GDH) and purified enzyme at 100 nM. Enzyme kinetic parameters (Km, Vmax, Vmax/Km and kcat) were determined by non-linear regression fitting of experimental data using Prism - GraphPad v7.00 software.

To evaluate the inhibitory effect of L-ascorbic acid on L-galactose dehydrogenase activity, a 200 μL reaction mixture containing 100 mM Tris-HCl pH 7.0, 200 μM of NAD^+^, L-gal (1 mM for *So*GDH and 2 mM for *Md*GDH) and 100 nM purified enzyme in the presence of varying AsA concentrations (0.75, 1.5, 3.0 and 4.5 mM) were monitored as described above. Likewise, the inhibition was also monitored using a more concentrated buffer (300 mM Tris-HCl pH 7.0). The percentage inhibition was calculated using the following equation: 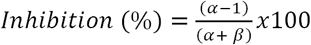, where: α is (1+[I])/*KI*, β is [S]/*Km*, [I] is inhibitor (α+*β*) concentration and [S] is substrate concentration.

### Protein crystallization

*So*GDH was crystallized by the sitting drop vapor diffusion method using the IndexTM HT screening kit (Hampton Research, Laguna Niguel, CA, USA). A drop of 0.2 μL of freshly purified protein (10 mg/mL) was mixed with 0.2 μL of the reservoir solution and suspended over the latter at 291 K. After 48 h, crystals were identified in the drop suspended over the reservoir solution consisting of 0.2 M Ammonium acetate, 0.1 M BIS-TRIS pH 5.5 and 25% (w/v) PEG 3350. Subsequently, crystals of *So*GDH complexed with the cofactor NAD^+^ were obtained using co-crystallization. In this case *So*GDH (10mg/mL) was mixed with 5 mM NAD^+^ and 1 mM L-gal in the same reservoir solution and crystallized in a similar fashion to that of the ligand-free enzyme. In all cases the crystals were harvested and cryo-cooled in liquid nitrogen for data collection.

### Data collection and structure determination

The X-ray diffraction data were collected at 100 K on the *Sirius* synchrotron (LNLS-CNPEM, Campinas) using beamline Macaná housing a PILATUS 2M detector for cofactor-free *So*GDH and on the *Diamond Light Source* synchrotron (Didcot, UK) using beamline I03 with an Eiger2 XE detector for *So*GDH-NAD^+^. The data were indexed, integrated, and scaled using Xia2 (Winter, 2010), and the structure of *So*GDH was solved by molecular replacement using *MoRDa* (Vagin and Lebedev, 2015). Subsequently, *So*GDH-NAD^+^ was solved by molecular replacement with the previously refined *So*GDH structure as the search model using Phaser (McCoy, 2007). Alternate rounds of refinement and model rebuilding conducted using Phenix (Adams et al., 2010) and Coot (Emsley and Cowtan, 2004) yielded the final models. The residues involved in the NAD^+^ binding site were determined with the Discovery Studio Visualizer V21.1.0 (BIOVIA, Dassault Systèmes) and figures were generated with PyMol v2.05 (Schrödinger, LLC). The data collection, refinement statistics, and PDB codes are summarized in Table S3.

### Protein structure prediction

*Md*GDH structure prediction was performed with ColabFold, an online extension of the AlphaFold2 software (Jumper et al., 2021; Mirdita et al., 2021). An MMseqs2 (Uniref+environmental) msa_mode, automatic model_type, unpaired+paired pair_mode and 3 rum_recycles were selected as the input parameters. One hundred and fifty sequences were selected as templates for model building, obtaining 5 final models. The quality of the modeled tridimensional protein structures was evaluated by IDDT parameters (Jumper et al., 2021; Mirdita et al., 2021) and stereochemistry by Procheck (Laskowski et al., 1993) and Verify3D (Eisenberg et al., 1997).

### Phylogenetics analysis

The evolutionary history of 13 subfamilies of AKRs annotated from the University of Pennsylvania PKR database and 5 representatives of plant L-GHDs was inferred using the Maximum likelihood method (Goldman, 1990). A bootstrap consensus tree was inferred from 1000 replicates (Felsenstein, 1985), with subfamily branches collapsed where possible. Branches corresponding to partitions reproduced in less than 30% bootstrap replicates were also collapsed. Phylogenetic analyses were conducted in MEGA11 (Tamura et al., 2021).

## Supporting information

Suppelmentary information

## Data Availability

The data underlying this article are available in the GenBank (accession number OK632632.1) and Protein Data Bank (PDB code 7SMI and 7SVQ).

## Acknowledgments

We gratefully acknowledge to Macromolecular Crystallography School “From data collection to structure refinement and beyond” - Montevideo (2021) for help in the collection and processing of the crystallographic data for *So*GDH-NAD^+^. We are also grateful to Susana Sculaccio and Andressa Alves Pinto for excellent technical support. We recognize the essential role played by DIAMOND–UK and CNPEM-BR by providing access to beamlines and to their technical staff during synchrotron data collection. We are grateful to Zui Fujimoto for depositing and releasing the coordinates for the rice GDH structures.

