## Supplementary material for "Structural Characterization of L-Galactose Dehydrogenase: A Key Enzyme for Vitamin C Biosynthesis": Suppelmentary information

**Supplementary information**

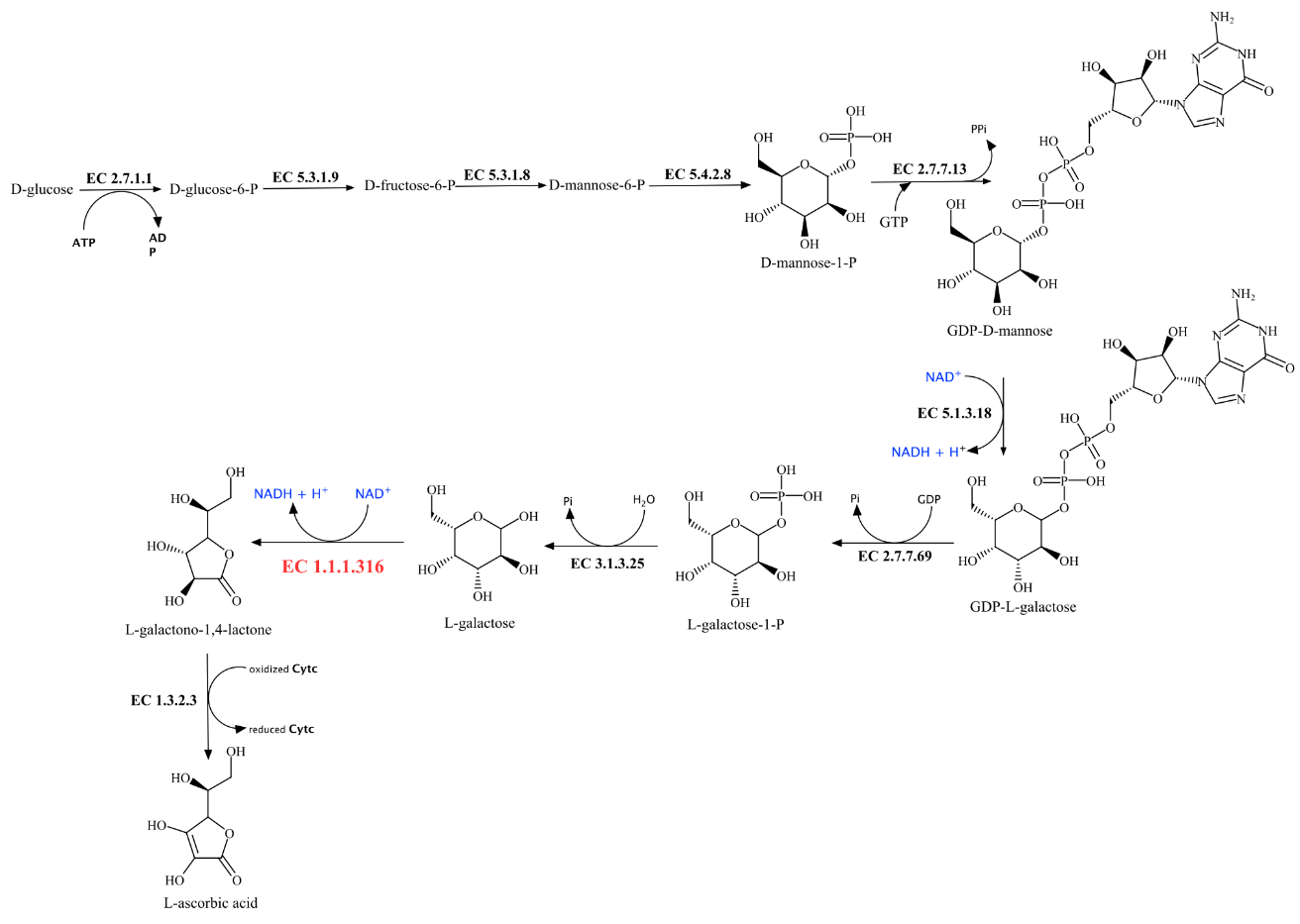

Figure S1. The D-mannose/L-galactose biosynthetic pathway for vitamin C. The step catalyzed by GDH (EC 1.1.1.316) is indicated in red.

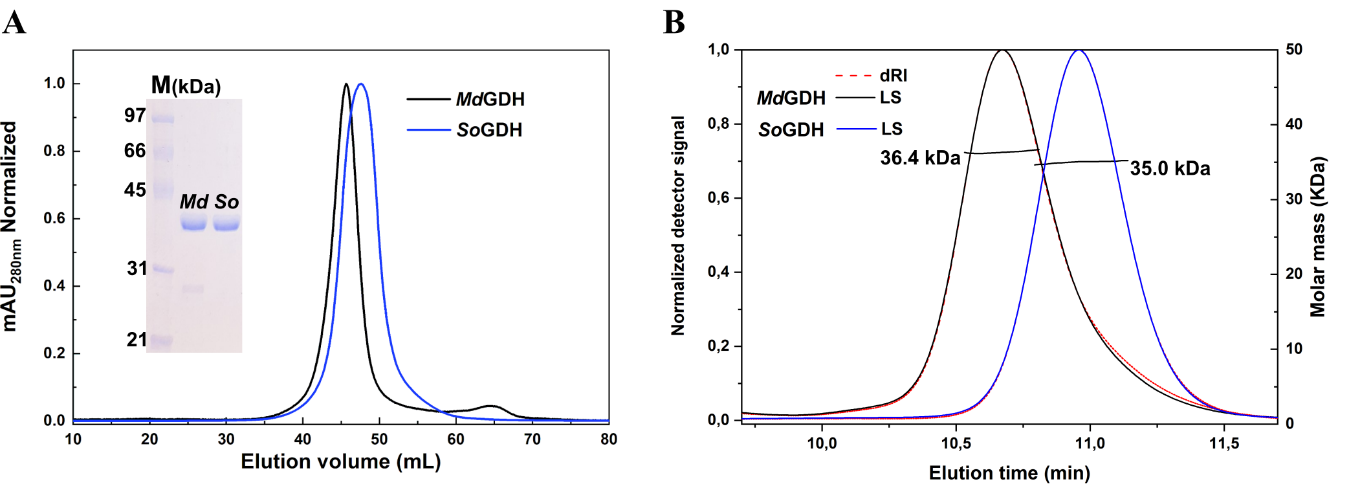

Figure S2. Recombinant MdGDH and SoGDH. (A) SEC profile with normalized absorbance_280nm_ signal. Both MdGDH and SoGDH show a predominant peak with a single band on SDS-PAGE, suggesting a single oligomeric state. (B) SEC-MALS traces. The curves corresponding to the change in the normalized differential refractive index (red), normalized scattered light intensity at 90º (black for MdGDH and blue for SoGDH) and calculated molecular weight of the corresponding peak (horizontal black line) are given. The calculated MW corresponds to the theoretical MW for a monomer of L-GDH.

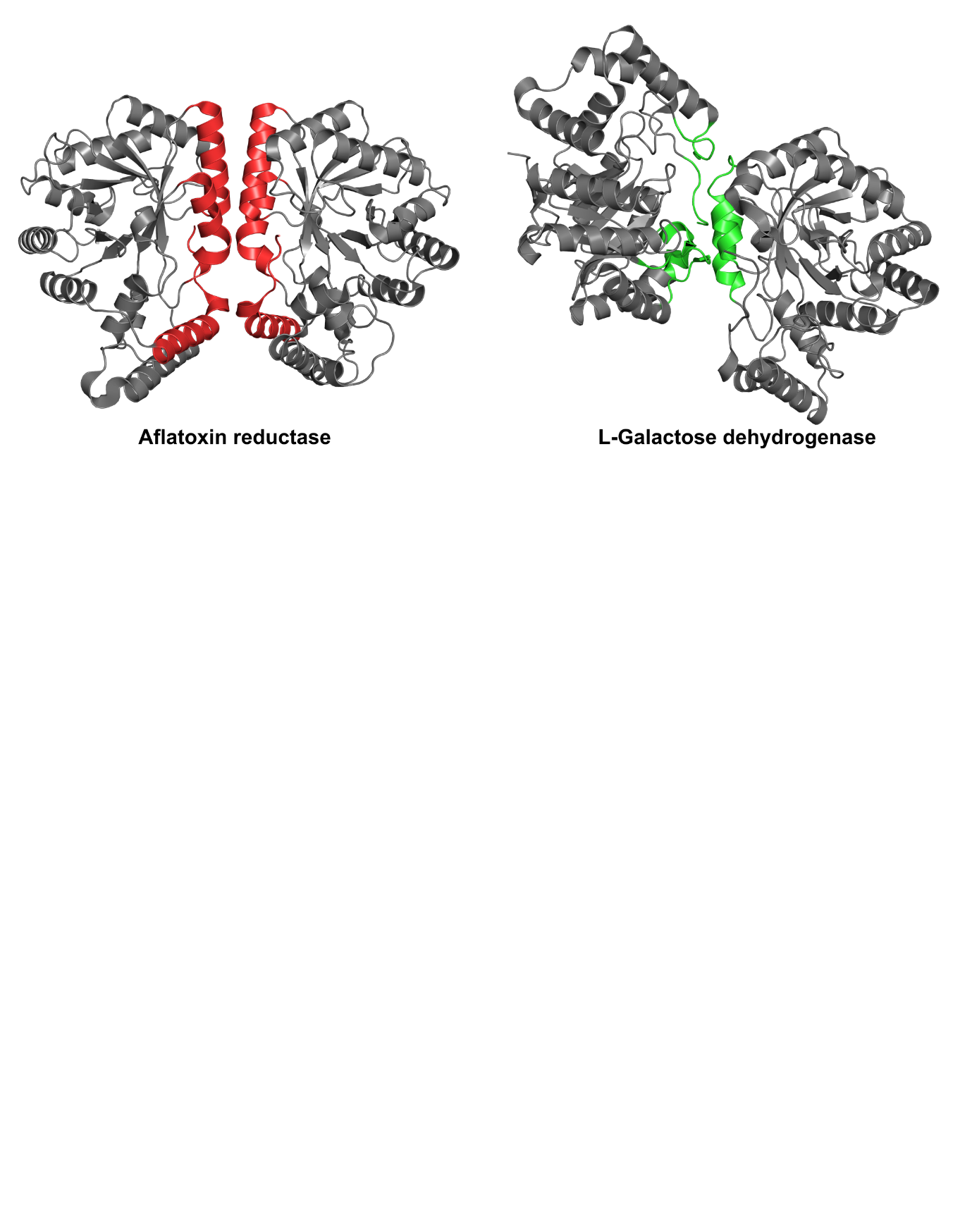

Figure S3. Dimerization interface in homodimeric AKRs (represented by aflatoxin reductase) compared with and the structure of *So*GDH-NAD^+^. Aflatoxin reductase uses α5, α6, H2 and Loop-C helices for dimerization (red). However, in the structure of *So*GDH-NAD^+^ the two monomers of the asymmetric unit make crystal contacts through loop β5-α3, Loop-B and Loop-C of chain A and the helices α5, α6 and loop α5-β8 of chain B (green). Aflatoxin reductase PDB code: 1GVE.

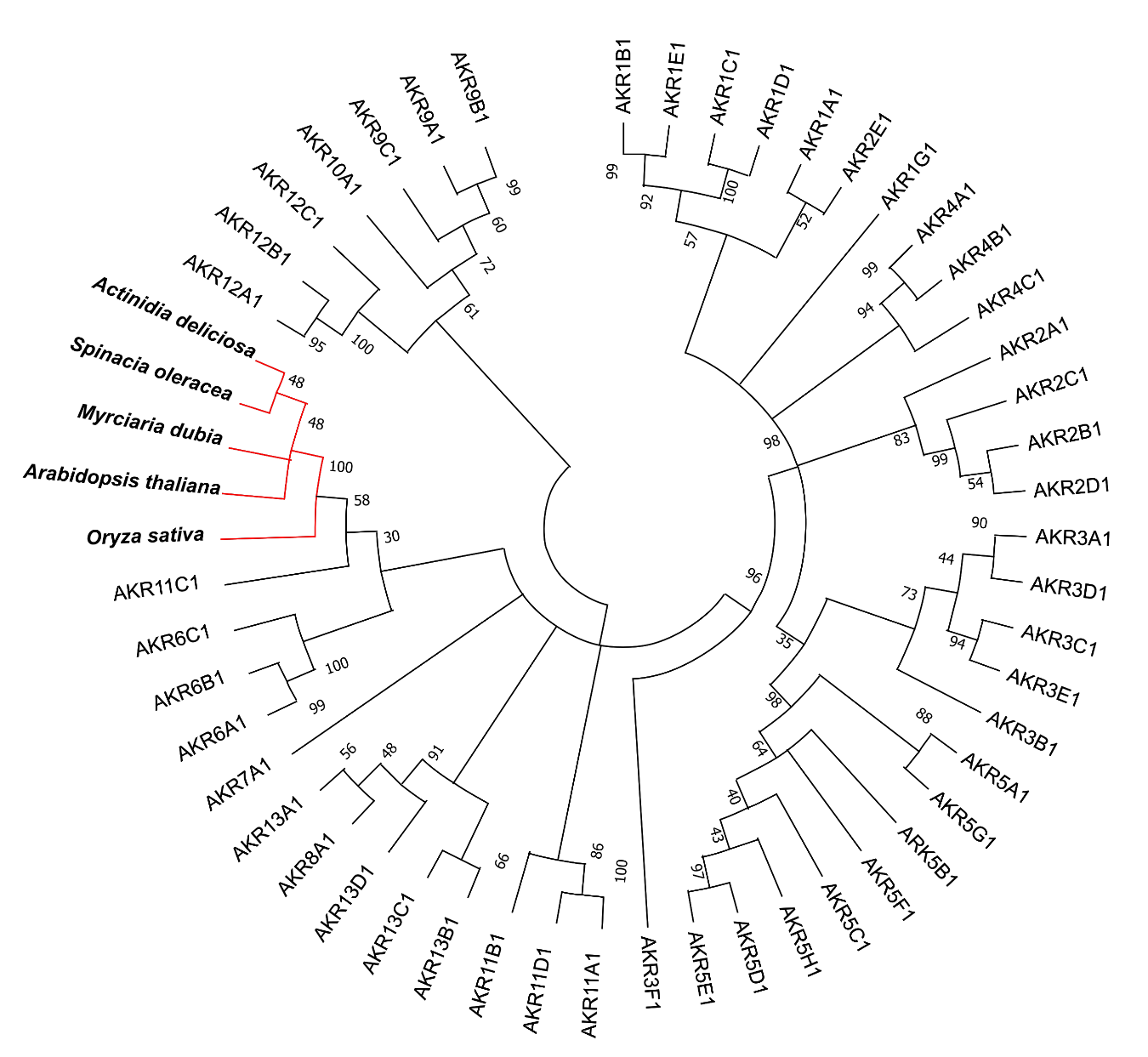

Figure S4. The dendrogram shows the topological relationships between some members of the AKRs and 5 representatives of L-GDH. The bootstrap consensus tree inferred from 1000 replicates shows L-GDHs to cluster into a monophyletic group (red branches), suggesting the formation of a new subgroup of AKRs by L-GDHs.

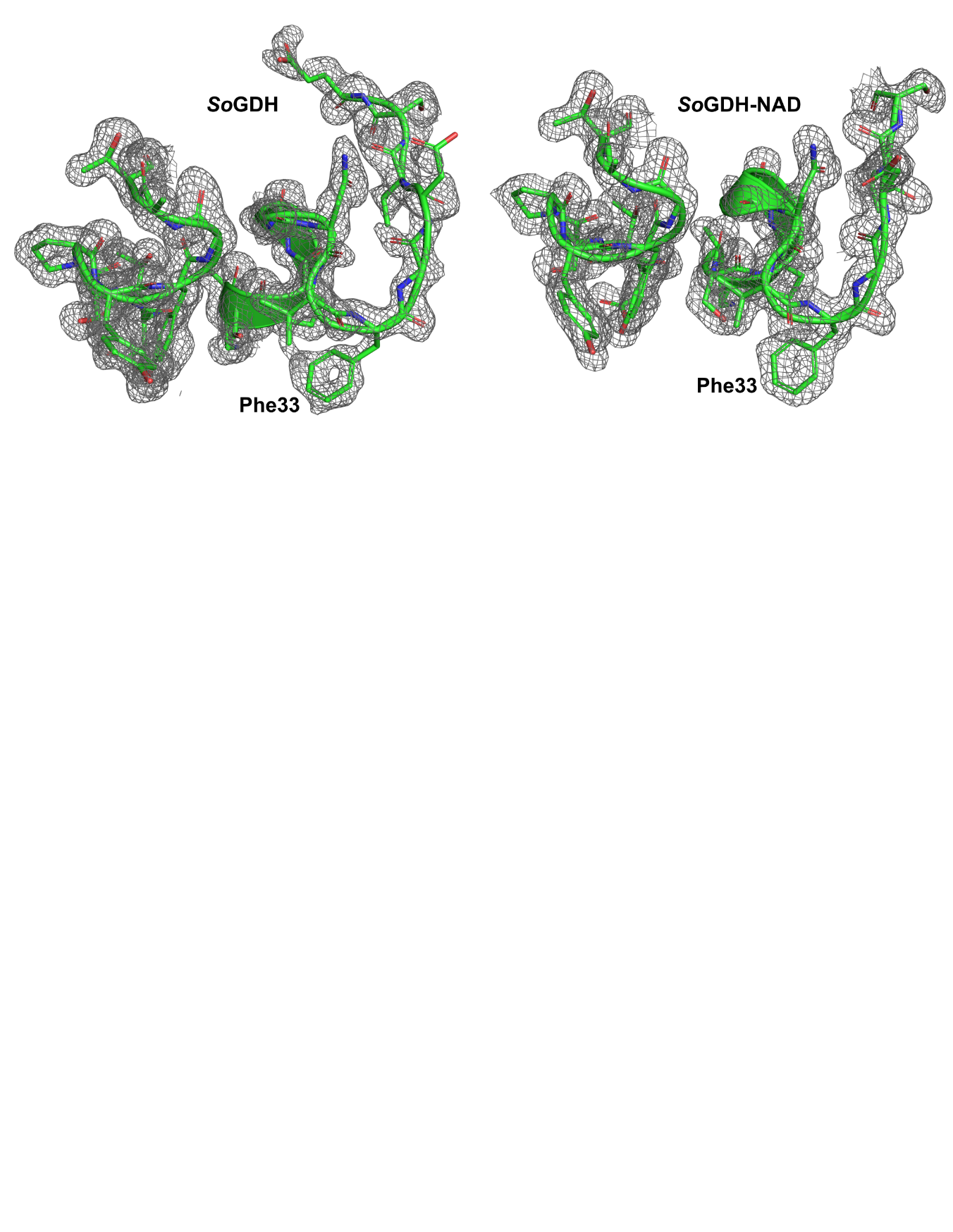

Figure S5. Electron density map (2Fo − FC at 1σ) of Loop-β3-α1 and Loop-β4-α2 in *So*GDH and *So*GDH-NAD^+^. The density clearly shows the orientation of Phe33 in both states.

**Table S1.** pH measurements in the reaction buffer

| AsA (mM) | 100 mM Tris-HCl | 300 mM Tris-HCl |
| --- | --- | --- |
| 0 | 7.02 | 7.11 |
| 0.75 | 6.87 | 7.10 |
| 1.5 | 6.78 | 7.09 |
| 3 | 6.60 | 7.03 |
| 4.5 | 5.90 | 6.99 |

**Table S2.** X-ray data collection and structure refinement statistics.

|  | *So*GDH | *So*GDH – *NAD^+^* |
| --- | --- | --- |
| X-ray source | Sirius MANACA | Diamond I03 |
| Detector | Pilatus 2M | Eiger2 XE |
| Cell parameters: a, b, c (Å)  Cell parameters: α, β, γ (°) | 51.72, 55.01, 61.79,  90.00, 112.10, 90.00 | 49.99, 54.97, 63.22  86.84, 69.13, 84.14 |
| Space group | *P*2_1_ | *P*1 |
| Resolution (Å) | 39.69 – 1.40 (1.42 – 1.40) | 39.73 – 1.75 (1.78 – 1.75) |
| λ (Å) | 1.3236 | 0.97622 |
| Multiplicity | 12.7 (12.1) | 3.4 (3.3) |
| *R*_pim_ (all I+ & I-) (%) | 4.0 (48.5) | 6.9 (38.1) |
| CC(1/2) | 0.998 (0.643) | 0.958 (0.706) |
| Completeness (%) | 99.6 (97.6) | 87.5 (40.8) |
| Reflections | 804980 (36548) | 190355 (4645) |
| Unique reflections | 63165 (3024) | 55241 (1416) |
| *<I*/*σ*(*I*)> | 14.0 (2.7) | 7.5 (1.8) |
| Reflections used in refinement |  |  |
| *R* (%)* | 19.42 | 18.37 |
| *R*_free_ (%)* | 20.95 | 21.48 |
| Number of atoms: protein | 2421 | 4845 |
| Number of atoms: water | 250 | 533 |
| Number of atoms: ligand | 0 | 88 |
| B (Å^2^) | 20.08 | 14.07 |
| Coordinate error (ML-based) (Å) | 0.14 | 0.19 |
| Phase error (°) | 21.96 | 24.50 |
| Ramachandran favored (%) | 97.76 | 98.08 |
| Ramachandran allowed (%) | 2.24 | 1.92 |
| All-atom clashscore | 1.45 | 2.87 |
| Bond lengths (RMSD) (Å)* | 0.009 | 0.006 |
| Bond angles (RMSD) (°)* | 1.043 | 0.790 |
| PDB entry | 7SMI | 7SVQ |

*Taken from depositor
